# Detection of inertial effects in capillary flows in open and closed channels

**DOI:** 10.1101/2025.02.24.639324

**Authors:** Jean Berthier, Jodie C. Tokihiro, Anika McManamen, TJ Caira, Albert Shin, Jing J. Lee, Ashleigh B. Theberge, Erwin Berthier

**Affiliations:** Department of Chemistry, University of Washington, Box 351700, Seattle, Washington 98195, United States; Department of Urology, School of Medicine, University of Washington Seattle, Washington 98105, United States

**Author notes:** Equal contribution (Jean Berthier, Jodie C. Tokihiro, Anika McManamen).

## Abstract

The Lucas-Washburn-Rideal law is commonly applied to describe capillary flow dynamics in closed or open channels, microporous media, such as paper pads and fiber threads or even granulous soil. It assumes a viscous flow regime where capillary forces are counteracted by friction with the solid structure, a valid assumption given the small flow velocities and device dimensions. However, scenarios exist outside the viscous regime, where inertial effects become significant, meaning capillary and friction forces do not fully balance. One well-documented case is the transient inertial regime at the onset of capillary motion. With the advancement of capillary devices, other configurations also raise the possibility of inertia influencing flow behavior. This study introduces a criterion to identify inertial contributions in capillary-driven flows in spatially varying geometries within open or closed channels and demonstrates how the Bosanquet equation can account for inertial effects in rectangular open-channel configurations.

## 1. Introduction

The Lucas-Washburn-Rideal (LWR) law [1-3]—and its modified form which takes into account complex non-circular geometries, denoted here mLWR [4,5]—is commonly used to describe capillary flow dynamics in both closed [6,7] and open channels [8-11], as well as in microporous media like paper pads and threads [12,13] and granular media such as soil [14]. It operates under the assumption of a viscous regime, where capillary forces are balanced by friction with the solid structure. However, there are situations beyond the viscous domain where inertial forces come into play. One well-known instance is the initial inertial regime at the beginning of capillary motion, where the wall friction surface area is small and the capillary force is much greater. This imbalance between capillary force and wall friction causes the flow to accelerate suddenly until friction balances with the capillary force [15-18]. This phenomenon, called initial inertial regime, occurs not only in tubes and closed or open channels but also in yarns and porous media [19].

Less commonly acknowledged are instances where the imbalance is caused by a sudden shift in capillary force. A characteristic case is that of an abrupt change in the channel’s geometry or a gradual variation in its cross-sectional area [20-23]. In these situations, during the morphological transition, the friction force may remain relatively unchanged on a short scale, but the capillary force—acting at the front meniscus—suddenly shifts, creating an imbalance between the two forces. This imbalance triggers an inertial effect, causing a perturbation of the flow velocity. In corroboration with pre-existing research, we refer to this as the viscous-inertial regime [23]. Most of the time this change of velocity is quickly damped by wall friction, but not always, as shown by Shobeiri et al. [23]. Another example, as we shall see, is the shift motion of a plug placed ahead of an open capillary flow [24]. It will be shown that the inertial effect is caused by a periodic oscillation of the capillary force.

In the initial phase of this study, we introduce a simple experimental nondimensional criterion to account for the potential appearance of inertial effects during capillary flow. The proposed criterion is derived by examining a characteristic function—denoted *F*—along the flow path, defined as the product of the cross-sectional area at the meniscus position, the travel distance, and the meniscus velocity—deduced as the time derivative of the travel distance. The function *F* is conveniently rendered dimensionless by dividing it by its value *F*_*0*_ at the device’s inlet channel when a viscous regime is established. By combining the geometric profile of the device with measurements of the travel distance, the presence of inertial effects can be identified. A constant domain of the function corresponds to a viscous regime. Oppositely, an increase or decrease of the function shows an inertial effect. We validate this approach by analyzing the dynamics of the capillary flow in open channels featuring converging and diverging sections, as well as bifurcations and also two-phase capillary flow.

In scenarios where inertial effects are absent or negligible, previous studies have demonstrated that the mLWR law accurately describes the flow dynamics in channels of arbitrary shapes [5,9,24,25]. This modified law incorporates a friction length to account for wall friction and a generalized Cassie angle to address different wall surface energies or the presence of a free surface in open capillary flows. Furthermore, at sufficiently high velocities, a dynamic contact angle (DCA), based on contact line friction [26,27], is included in the formulation. Evaluating the applicability of the mLWR requires identifying the presence and significance of inertial effects. When these effects become significant, the Bosanquet law—or, for open channels, the modified Bosanquet law (mBo)—must be applied. The mBo law also includes a friction length, a generalized Cassie angle, and a dynamic contact angle (DCA), akin to the mLWR. Since inertia is typically associated with higher velocities, the inclusion of the DCA is usually essential when using the Bosanquet law.

Here, we find that inertial effects are not uncommon in capillary devices, but the geometric variations must have sufficient spatial extent to significantly influence the overall dynamics. This explains the continued applicability of the mLWR law in many cases. Nonetheless, when substantial inertial effects occur due to the spatially varying channel geometry, it becomes necessary to utilize the Bosanquet equation [24,28,29], which incorporates an inertial term.

In the second part of this study, we analyze flow dynamics using the mBo law for converging and diverging open rectangular channels, validating the results through experiments conducted with 20% (v/v) aqueous isopropyl alcohol (IPA 20%), 50% (v/v) aqueous isopropyl alcohol (IPA 50%), and pentanol. The influence of inertia is highlighted by comparing the Bosanquet approach to the non-inertial solution, using experimental data from rectangular open channels fabricated in poly(methyl methacrylate) (PMMA). It is shown that a viscous-inertial motion takes place in convergent channels, while the inertial effect is small or negligible in divergent open-channels. Additionally, we observe that flow velocities increase in converging channels and decrease in diverging channels. Although our analysis focuses on open channels, it can be readily extended to closed-channel systems.

## 2. A criterion for inertial effects

In the LWR approach for the viscous capillary regime, the capillary and wall friction forces balance each other, i.e., *F*_*cap*_ = *F*_*drag*_. This approach was has been successfully used for bifurcating channels [30,31], capillary trees [32,33], etc. Note that the LWR law results in a relation between the travel distance *z* and velocity *V* of the form *z V* = *cste*, under the assumption of a static contact angle [27]. For capillary flows in open channels, the LWR law must be modified [5] by introducing a friction length to account for the influence of the free surface and the non-circular channel geometry. Additionally, a generalized Cassie contact angle is incorporated to average the effects of the various walls and the free surface [4].

On the other hand, for a viscous-inertial regime, the Bosanquet equation [28] can be written as

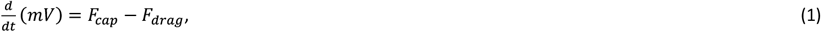

where *m* is the mass of the moving liquid, *V* its velocity, and *F*_*cap*_and *F*_*drag*_the capillary and friction force. If we take the example of an open-channel capillary flow with a varying channel width *w* (and a constant height *h*) as sketched in Figure 1, and use the mass conservation equation between any location in the channel and the location of the front meniscus (referred by the index *f*): *w*_*f*_ *V*_*f*_ = *w*(*z*) *V*(*z*), the inertial term becomes

**Figure 1.**
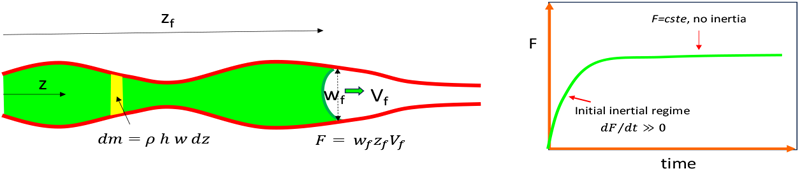
Left: sketch of the open channel with varying width *w;* right: an inclined function F corresponds to an inertial situation while a horizontal (constant) function F corresponds to a no-inertia case.

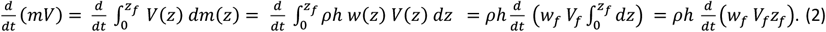

where *ρ* is the liquid density, *t* the time, *z* the axial coordinate, *V* the velocity in a cross section, and *z*_*f*_ the location of the flow front.

Hence, the variation of the term *F* = *h w*_*f*_ *V*_*f*_*z*_*f*_ —uniquely corresponding to the characteristics of the front of the flow (advancing meniscus)—indicates the occurrence of inertial effects. In the case where the depth *h* is not constant, we can incorporate the height *h* in the function *F* to account for a change in the channel depth, so that

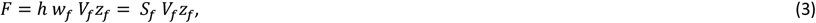

where *S*_*f*_ is the cross-sectional area. In what follows, we found it convenient to nondimensionalize the function *F* by dividing it by the constant inlet value *F*_*o*_ obtained when the advancing meniscus travels through the inlet channel, so that *F*/*F*_0_ = *w*_*f*_ *V*_*f*_*z*_*f*_/*w*_0_ *V*_0_*z*_0_ (*inlet*).

So, on a plot of the function *F*, the horizontal parts correspond to non-inertial regime, i.e., viscous regime, while the sloping regions correspond to the presence of inertial effects. In the following sections, we illustrate this principle by examining experimental results.

## 3. Experimental examples of inertial and non-inertial capillary flows

We analyze five different experiments. For the first two (convergent and divergent channels), we designed and performed the experiments in this study. For the last three (bifurcating channels[31], capillary tree[32,33] and two-phase flow[34]), we utilized data or methods from previously published experiments.

### 3.1 Example #1: convergent open-channels

The first example corresponds to open devices with a linearly converging section, as shown in Figure 2A, where the channel is an open rectangular channel milled in PMMA. The dynamics of liquids such as nonanol and IPA 50% have been monitored by locating the travel distance (location of the front meniscus) with time, *z(t)* as shown in Figure 2B. Travel velocity is then deduced by time derivation: *V*_*f*_ = *dz*_*f*_/*dt* (Figure 2C). The inertial function *F* = *h w*_*f*_ *V*_*f*_*z*_*f*_ is finally derived (Figure 2D), showing the expected inertial regime in the inlet and an inertial effect in the convergent. The inertia force *I* = *d*(*mV*)/*dt* = *dF*/*dt* is strongly positive at the inlet, positive in the convergent, and negative in the convergent exit. Hence, at the convergent entrance, the capillary force becomes larger than the wall friction along the whole device, and at the convergent exit, the capillary force is less than the wall friction.

**Figure 2.**
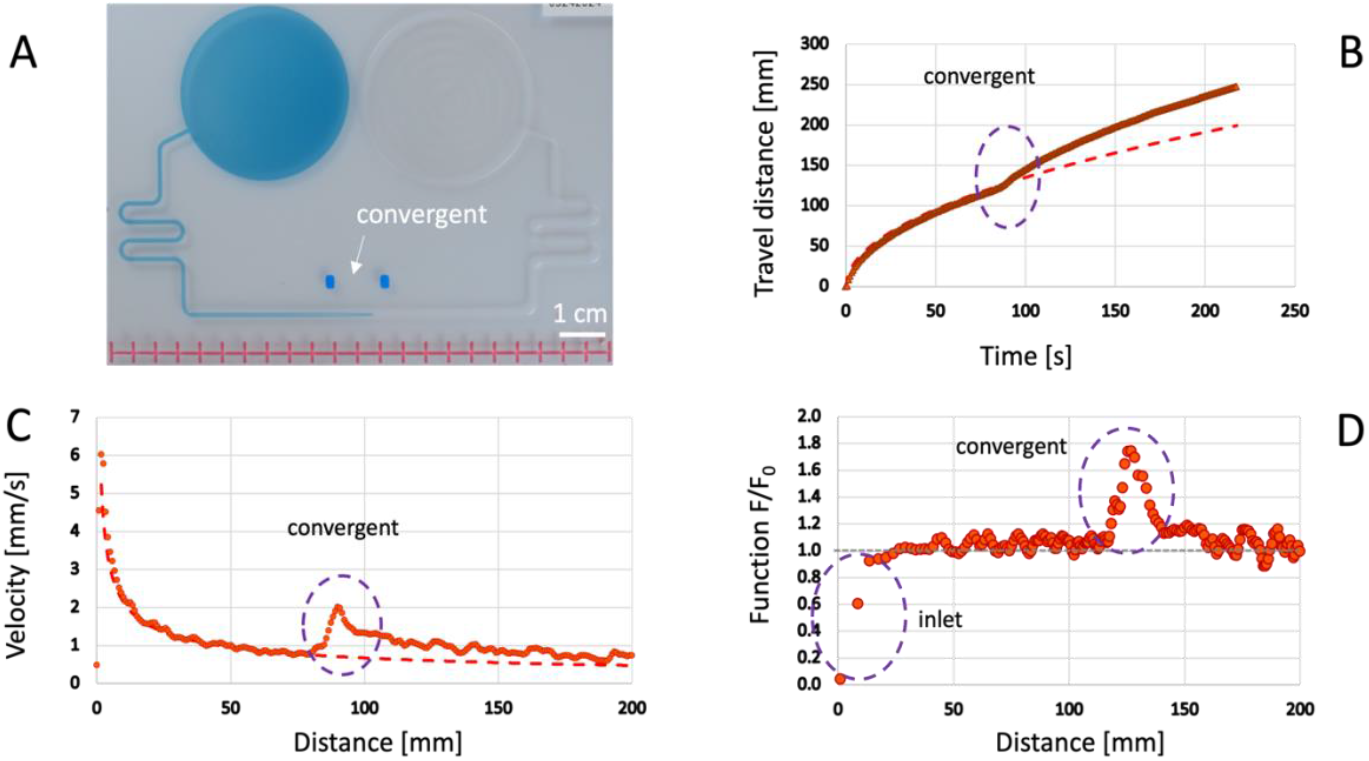
A: perspective view of the device with colored nonanol flowing in an open-rectangular 1 mm x 0.5mm channel reducing to 0.5 mm x 0.5 mm channel along a convergent distance of 10 mm; B: measured travel distances vs. time, the dashed line represents the mLWR law as applied to an extended inlet channel, thereby highlighting the influence of the converging section; C: velocities vs. distance, showing a velocity increase in the convergent channel; D: non-dimensional inertial function *F*/*F*_0_, showing a strong inertial effect at the beginning of the capillary flow where capillary force is much larger than wall friction, and a sharp peak during the passage of nonanol in the converging channel. This peak is much higher than the fluctuations of the function due to measurement errors. Data are reported as an average of three replicates (n = 3).

### 3.2 Example #2: divergent open-channels

The second set of experiments resembles the previous ones; however, this time, the channel includes a linearly divergent section (Figure 3). The function *F/F*_0_ for this case is plotted in Figure 3. The inertial motion at the channel entrance is clearly visible, while the divergent section introduces only a minor or negligible inertial effect, depending on the liquid properties and the dimensions of the divergence.

**Figure 3.**
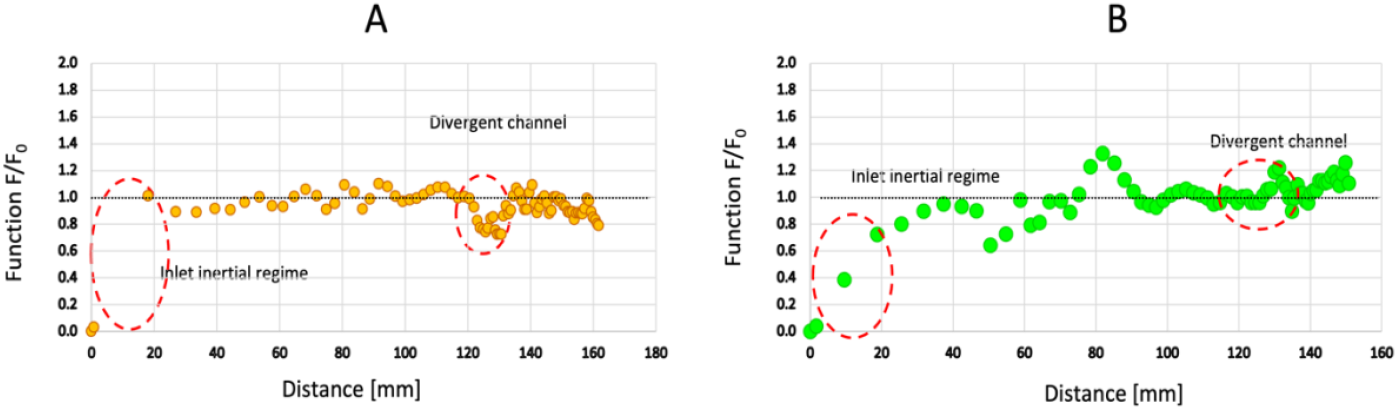
A: non-dimensional inertial function *F*/*F*_0_ = *w*_*f*_ *V*_*f*_*z*_*f*_/*F*_0_ for an IPA 50% v/v showing a small inertial effect in the divergent (length of divergent 10 mm, width at entrance 0.5 mm, width at exit 1 mm, channel depth 0.5 mm); B: non-dimensional function *F/F*_*0*_ for pentanol flow in a divergent open-channel showing no inertial effect (length of divergent 20 mm, width at entrance 1 mm, width at exit 2 mm, channel depth 2 mm). Some fluctuations of *F/F*_*0*_ up to 30% during the motion in the root channel and exit channel can be seen on the graph due to travel distance measurement errors and adjustment of the liquid level in the inlet reservoir (in order to maintain a zero pressure). Data are reported as an average of three replicates (n = 3).

### 3.3. Example #3: bifurcations

Closed-channel bifurcations were analyzed by Mehrabian et al. for both symmetric and asymmetric cases [30], while Jing et al. investigated similar bifurcations in open channels [31], arriving at comparable findings. In symmetric bifurcation, the flow velocity remains identical in both daughter branches. However, in asymmetric bifurcation, the velocity is initially larger in the smaller channel before declining after a certain distance. This phenomenon results from the interplay between capillary forces and friction in both channels.

For our third example, we utilize velocity data derived from [31], where open rounded-bottom rectangular channels divide into two daughter channels with the same shape, either symmetrically or asymmetrically, as depicted in Figure 4. The channel dimensions are detailed in the figure captions. Notably, it was established that the mLWR law accurately predicts the velocities in both daughter channels, indicating that significant inertial effects are absent in these experiments. Let us examine the situation using the inertial criterion: before the bifurcation, *F* is given by *F*_0_ = *h* _*f*0_*w*_*f*0_ *V*_*f*0_*z*_*f*0_, while after the bifurcation it is expressed as *F*_12_ = *h* _*f*1_*w*_*f*1_ *V*_*f*1_*z*_*f*1_ + *h* _*f*2_*w*_*f*2_ *V*_*f*2_*z*_*f*2_. In the figure, we have plotted the nondimensionalized function *F*. Whereas the plot of the function *F* does not show any particular effect at the symmetric bifurcation (Figure 4A), a small peak is observed in the asymmetrical bifurcation. This behavior can be linked to the reduction of cross-sectional area of the smaller daughter channel in the asymmetric case.

**Figure 4.**
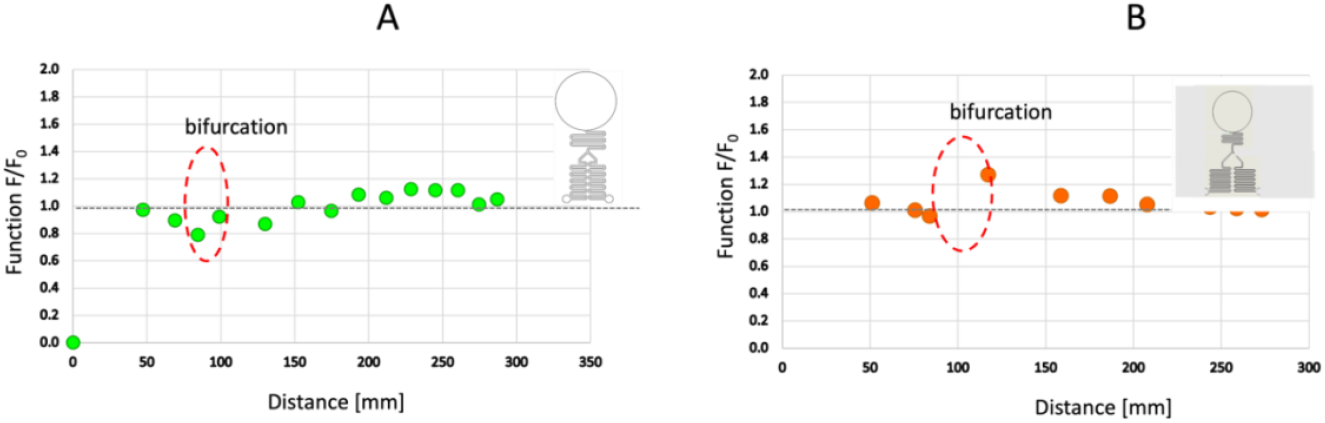
Two examples of open flow of nonanol through a symmetrical and asymmetrical bifurcation: A, symmetric bifurcation, all cross-sections are identical (rectangular with rounded bottom, w=0.8 mm, h_max_=1.5 mm), the velocities are identical in the two daughter channels; B: asymmetrical bifurcation, root and daughter #1 cross-sections have a width of 0.8 mm and maximum depth 1.5 mm, daughter channel #2 has a cross section of width 0.4 mm and maximum depth 0.75 mm. Details of experiments are reported in [31]; data are reported as an average of three replicates (n = 3).

### 3.4 Example #4: capillary trees

The fourth example also illustrates a case of channel division and is based on references [32,33], which explore the flow dynamics in capillary trees (Figure 5A). In these experiments, successive bifurcations, each accompanied by a reduction in cross-sectional area, are used to achieve capillary pumping. Specifically, the cross-sectional area decreases homothetically by a factor of 0.8 or 0.85 after each bifurcation. The flow rates briefly increase after each bifurcation so that the flow rate in the root channel, which is the sum of the flow rates in the tree (*S*_0_ *V*_0_ = ∑_*i*_ *S*_*i*0_ *V*_*i*_) stays relatively high—justifying the term “capillary pumping.” The function *F*, plotted in Figure 5B for a nonanol flow [31] and 5C for an IPA 50% flow [32] remains approximately constant throughout the tree, except for small increases at the bifurcations. This suggests that inertial effects at the bifurcations are not important. This remark is justified by the adequation of the mLWR model documented in the literature.

**Figure 5.**
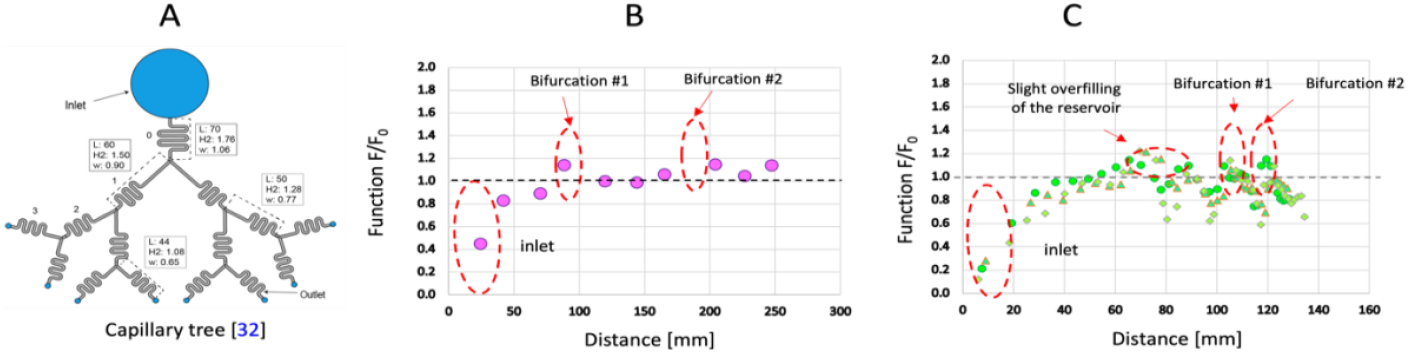
A: schema of the capillary tree milled in PMMA [31]; B: nonanol flow: at some distance of the inlet, the inertial function *F* is approximately constant—except for a small increase indicated in red on the figure. Data are shown for 1 replicate (n = 1); C: IPA 50% flow: at some distance of the inlet, the inertial function *F* is approximately constant—except for a small increase indicated in red on the figure. Details of experiments are reported in [31,32]. Data are shown for three replicates (n = 3). Reprinted (adapted) with permission from [34] J.J. Lee, J. Berthier, K.A. Brakke, A.M. Dostie, A.B. Theberge, E. Berthier, 2018 Droplet Behavior in Open Biphasic Microfluidics *Langmuir*, **34**, 5358-5366. Copyright 2025 American Chemical Society.

### 3.5 Example #5: two-phase capillary flows

The dynamics of a two-phase capillary flow of pentanol shifting ahead a plug of water in a rectangular open channel is our fifth example (Figure 6A). In this case, a 3 µl yellow water (10% yellow food dye) plug was loaded at a distance of 199 mm from the entrance into the channel (w = 0.8 mm, h = 1.5 mm), and 950 µL of blue pentanol (1 mm/µL Solvent Green III) was pipetted into the inlet reservoir. Sufficient blue pentanol was added to the reservoir to achieve a flat meniscus and prevent pressure-driven flow. As the meniscus formed and the plug moved, blue pentanol was slowly pipetted to maintain the flat meniscus until the pipette was emptied. Periodic velocity oscillations of the front end are observed (Figure 6B): when the carrier-pentanol front meniscus contacts the water plug, a sudden forward capillary force is exerted on the plug. The plug is submitted to an inertial regime with fast velocity, similar to that of a liquid entering a channel. The carrier liquid is, on the other hand, still flowing in the slow viscous regime. Hence, the plug tends to separate—at least partially—from the carrier liquid, and the resultant of the capillary force exerted on the plug rapidly decreases. These fluctuations of the capillary force trigger flow velocity oscillations that are superposed to the viscous regime velocity, as shown in Figure 6D. These oscillations are also seen in the plot of the inertial criterion function *F* (Figure 6E). The situation is typically a superposition of a basis viscous regime with inertial fluctuations. The “basis” line is defined by the viscous regime dynamics for pentanol (no plug), i.e. the relation 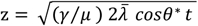 stated by the mLWR law (see table 1 for the notations and physical values).

**Table 1.** Physical properties of solvents.

|  | Density | Surface tension | Viscosity | Contact angle | Triple line friction coefficient |
| --- | --- | --- | --- | --- | --- |
| Units | [kg/m<sup>3</sup>] | [mN/m] | [mPA.s] | [degree] | [Pa.s] |
| Notation | $\rho$ | $\gamma$ | $\mu$ | $\theta$ | $\zeta$ |
| IPA 20% | 950 | 29 | 3.0 | 28 | 0.08 |
| IPA 50% | 900 | 31 | 3.2 | 34 | 0.10 |
| Pentanol | 814 | 27 | 3.7 | 23 | 0.05 |
| Nonanol | 827 | 28 | 11.5 | 48 | 0.10 |

**Figure 6.**
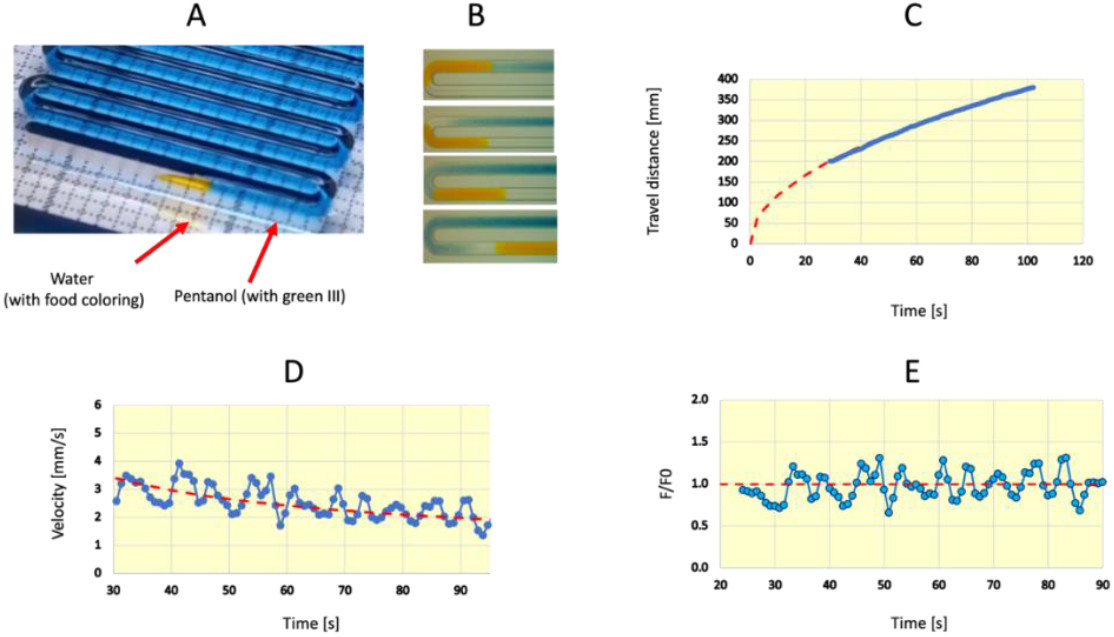
A: photograph of the pentanol (blue-green color) and water plug (yellow color) in the winding open channel; B: Four steps of the motion, showing pentanol contacting the plug, then the inertial “jump” of the plug, followed by a partial plug-pentanol separation, and finally a new contact of the pentanol with the plug; C: travel distance vs. time, the blue dots correspond to the experimental results and the discontinuous red curve to the “underlying” viscous regime. This regime corresponds to the relation 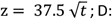 velocity vs. time showing the oscillations on each side of the “underlying” viscous regime; E: function *F/F*_*0*_ vs. time showing oscillations on each side of the horizontal “underlying” viscous regime; these oscillations are roughly periodic with a period of 6 seconds. Data shown are resultant of one replicate (n = 1).

### 3.6 Observations based on experimental results

The function *F* serves as a qualitative criterion for identifying inertial appearance in capillary flows. However, it is inherently noisy due to measurement errors in the travel distance *z*, which propagate further when determining the velocity *V* and become even more pronounced when calculating the product *zV*. However, an evaluation of the inertia criterion *F* in various experiments conducted in rectangular open channels reveals that inertia often becomes significant when there is a sudden change in capillary force. Such changes can arise from variations in the cross-sectional area or the presence of a diphasic flow. In cases where the cross-section undergoes local changes, the overall impact of inertia on flow dynamics remains minimal, as velocity jumps are quickly dampened by wall friction. The mLWR law for the viscous regime has been shown to accurately describe flow dynamics in these situations.

In contrast, we demonstrate that when the cross-sectional variation extends over a certain geometric length, as observed in convergent channels, this law no longer holds. In the next section, we will verify this statement through a detailed analysis of velocity and the inertia criterion function *F* in both linearly convergent and divergent channels.

## 4. Theoretical approach for a rectangular channel with converging section

The notations for the theoretical analysis and the physical properties of the liquids that have been used are listed in Table 1.

The open-rectangular channel is particularly interesting because these types of channels are commonly used in microfluidic circuitry. One of the main reasons is the facility of fabrication. The flow dynamics in such channels— in presence or not of capillary filaments [35-39]—have been well-documented when the channel cross-section remains constant [9,24,25,40] However, the flow dynamics become significantly more complex when the cross-section varies [41,42].

In this section, we apply the Bosanquet approach to incorporate the effects of inertia in a linearly converging section, as illustrated in Figure 2.

It has been shown that the capillary force in a uniform monolithic channel was (4)

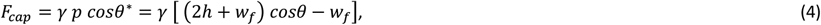

where *γ* is the surface tension, *p* the total channel perimeter in a cross section (free and wetted), and *θ*^∗^ the generalized Cassie angle [16]. A correction to the contact angle can be done by considering the dynamic contact angle but we do not show it here for simplicity. We just recall that, in the case of open-channels, it has been demonstrated that the expression of the dynamic contact angle (DCA), denoted *θ*_*d*_, was given by considering the triple contact line friction ξ [26,27],

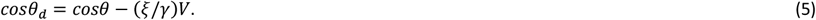

Substituting the expression (5) in (4) shows that the capillary force is not exactly constant in a uniform cross-section channel. In the following developments, we use the notation *θ* for simplicity instead of *θ*_*d*_.

In a linearly converging channel with a small geometrical angle *2α*, the capillary force is approximated by [24]

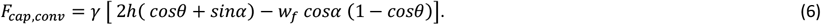

This expression is derived from Gibbs energy considerations [43] valid for small to moderate velocities. We have extended the use of the formula to this case where there is a small inertial effect. One checks that (6) reduces to (4) when *α*=0.

There are different approaches to determine the wall friction in open capillary flows. One of these is the decomposition in Fourier series of the velocity in the Stokes equation [9]. Another convenient approach is that of the friction length as proposed by Berthier et al. [5,24,25]. Let us recall that the friction length, denoted as 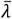, is a purely geometric length that represents the friction within a cross-section, where 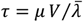. The wall friction in the convergent channel, between the distance *z=0* at entrance and *z=z*_*f*_ at the front end, is then given by:

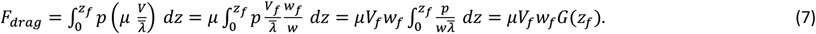

In (7), we have again used the mass conservation equation *w V* = *w V*. The integrand 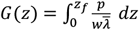 is a geometrical value characteristic of the shape of the channel. For example, for a rectangular channel of aspect ratio close to 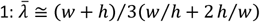 and *p* = *w* + *h*, hence

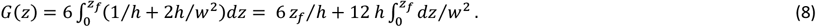

Then, when the flow front is located in the convergent, the Bosanquet equation (1) becomes

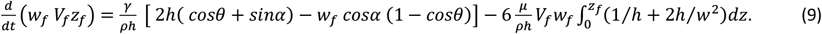

In the case of a linear convergent— the width *w*_*f*_ is a linear function of the distance: *w*_*f*_ = *χ*_1_ + *χ*_2_ *z*_*f*_. In such a case, the last term on the RHS can be integrated and equation (9) becomes the differential equation in *z*_*f*_:

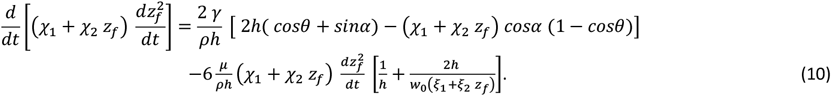

Note that the contact angle *θ* may depend on the velocity as stated in equation (4).

## 5. Results for converging channels

The solution of equation (10) is obtained by explicit discretization using the MATLAB (https://www.mathworks.com) software and is detailed in SI.1.

### 5.1. Experimental device

We created two rectangular open microfluidic channels with a geometrically converging cross-section going from dimensions (channel width x channel height) of (1) 1 mm x 0.5 mm to 0.5 mm x 0.5 mm and (2) 2 mm x 2 mm to 1 mm x 2 mm (Figure 7, engineering drawings in SI.3). The converging region progressively decreases in size only in the width dimension along a 10 mm length or from 10 mm to 30 mm for the 1 mm x 0.5 mm to 0.5 mm x 0.5 mm and the 2 mm x 2 mm to 1 mm x 2 mm devices, respectively. The device can reversibly be used for the study of the flow in convergent or divergent channels. The devices are also comprised of an inlet reservoir in which the liquid is added via a pipet, two long winding channels on each side of a convergent section, and an outlet reservoir at the extremity of the channels. The use of winding channels is required by the dimensions of the solid plate. Previous studies have shown that the turns do not affect the capillary flow in the absence of capillary filaments, as is the case in our study. The channels fabricated in PMMA and have rounded inner corners to minimize the formation of capillary filaments. The inlet and outlet reservoirs were made large to maintain constant Laplace pressure and markers were designed in the device to indicate the beginning and end of the converging region. Reservoirs were refilled once the fluid front exited the converging region. Using these devices, we flowed fluids of varying viscosities (e.g., IPA 50%, pentanol, nonanol) in converging and/or diverging configurations with physical properties listed in Table 1. The fluid dynamics in different experiments were compared between the inertial (mBo) and non-inertial approaches (mLWR).

**Figure 7.**
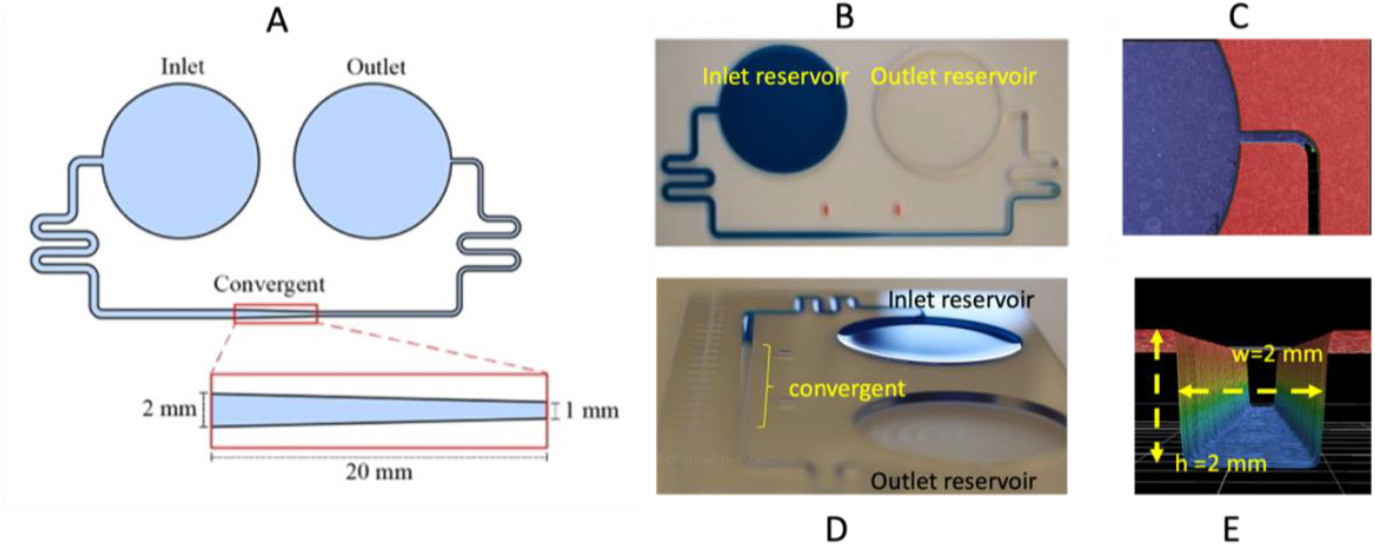
A: Schematic view of the device; B: View of the liquid motion through the device, the two red marks indicate the location of the convergent channel; C: View of the channel entrance with the profilometer; D: perspective image of the liquid entering the convergent; E: profilometer view of the channel cross section.

### 5.2. Results

Figures 8 shows a comparison between the measured travel distances and velocities and the results of the model for IPA 50% and pentanol in different converging channels. The plot of the travel distances shows an inflection at the passage of the convergent, and the plot of the velocities shows a local increase in the convergent. The velocity rebound is larger for shorter convergent in relation with a higher value of the convergent geometric angle *α*.

**Figure 8.**
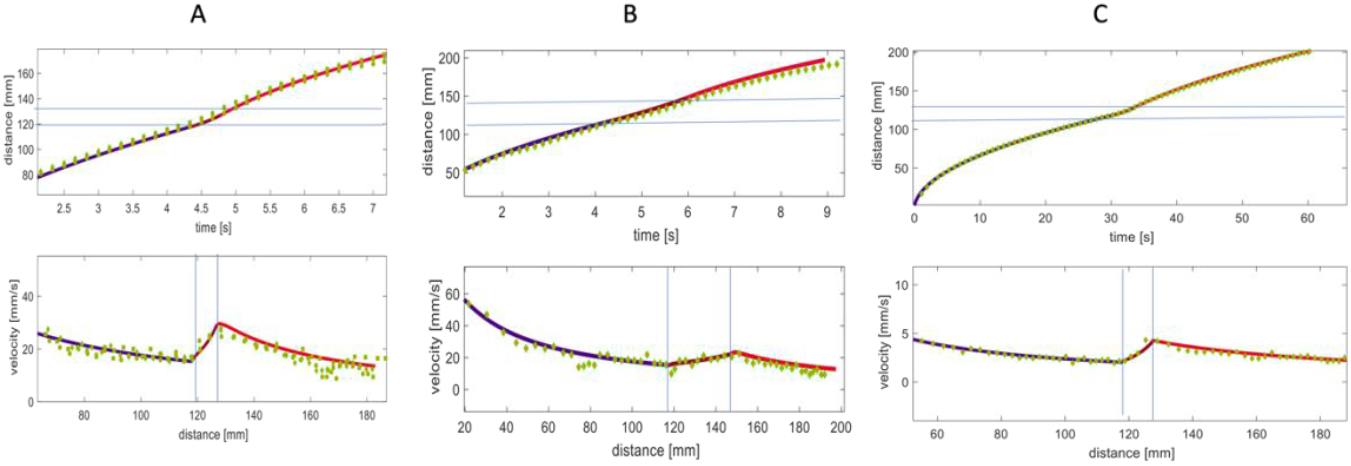
Comparison between experiments and model for (A) an IPA 50% flow in a 2 mm x 2 mm channel reducing to 1 mm x 2 mm in a 10 mm long convergent; (B) an IPA 50% flow in a 2 mm x 2 mm channel reducing to 1 mm x 2 mm in a 30 mm long convergent; (C) an IPA 50% flow in a 1 mm x 1 mm channel reducing to 0.5 mm x 1 mm in a 10 mm long convergent. The dots represent the experimental observations and the continuous lines represent the model predictions. The horizontal (top) and vertical (bottom) straight lines indicate the boundaries of the convergent. Data are reported as one trial (n = 1).

The inertia function *F* = *h w*_*f*_ *V*_*f*_*z*_*f*_ and the flow rate *Q* = *h w*_*f*_ *V*_*f*_ are plotted in Figure 9 for the case of a capillary flow of IPA 50% in a 10 mm linearly converging channel. The theoretical results (continuous line) coincide well with the experiments (dots). Note, the flow rate stays approximately constant in the convergent while it steadily decreases in the uniform cross-section channels upstream and downstream.

**Figure 9.**
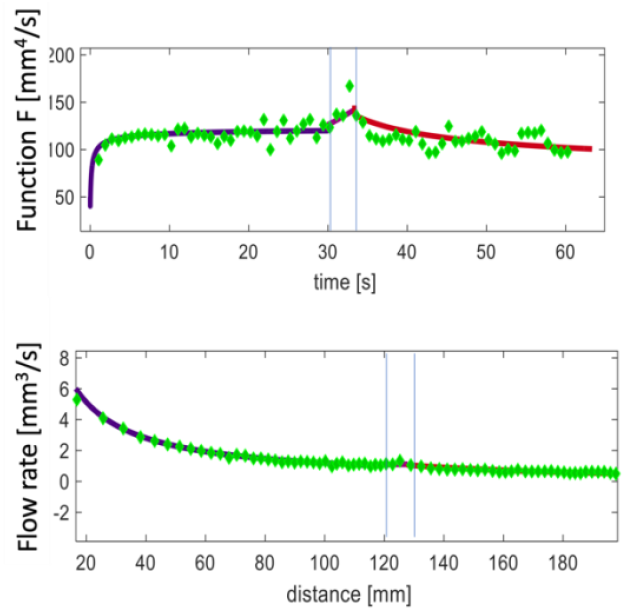
IPA 50% flow in a 1 mm x 1 mm channel reducing to 0.5 mm x 1 mm in a 10 mm convergent. The dots correspond to the experimental observations and the continuous lines to the results of the model. The vertical straight lines indicate the boundaries of the convergent. Top: inertia function *F* in function of time. Bottom: flow rate in the channel vs distance. Data are reported as one trial (n = 1).

### 5.3. Discussion: Comparison with non-inertial solution

Few investigations on the capillary-driven flow in converging channels have been reported. Gorce et al. studied the acceleration of the velocity in an axisymmetric convergent channel [20] and, in his Master Thesis, Zakershobeiri [22,23] indicates a numerical approach for closed channels with an axisymmetric shape that takes into account inertial effects. In our case of a rectangular open-channel, we compare the results of the mBo law to the mLWR law using a pressure approach, i.e., balance the local capillary pressure (Laplace pressure) with the total pressure drop along the walls (see SI2).

Figure 10 shows the comparison between the inertial model (mBo), the viscous model (mLWR), and experiments in the case of IPA 50% flowing in a 10 mm long converging channel and in the case of pentanol flowing in a 20 mm long open-channel. In both scenarios, a dynamic contact angle was employed. The viscous model exhibits a noticeable overshoot when compared to the experimental data, while the viscous-inertial model aligns well with the experimental results.

**Figure 10.**
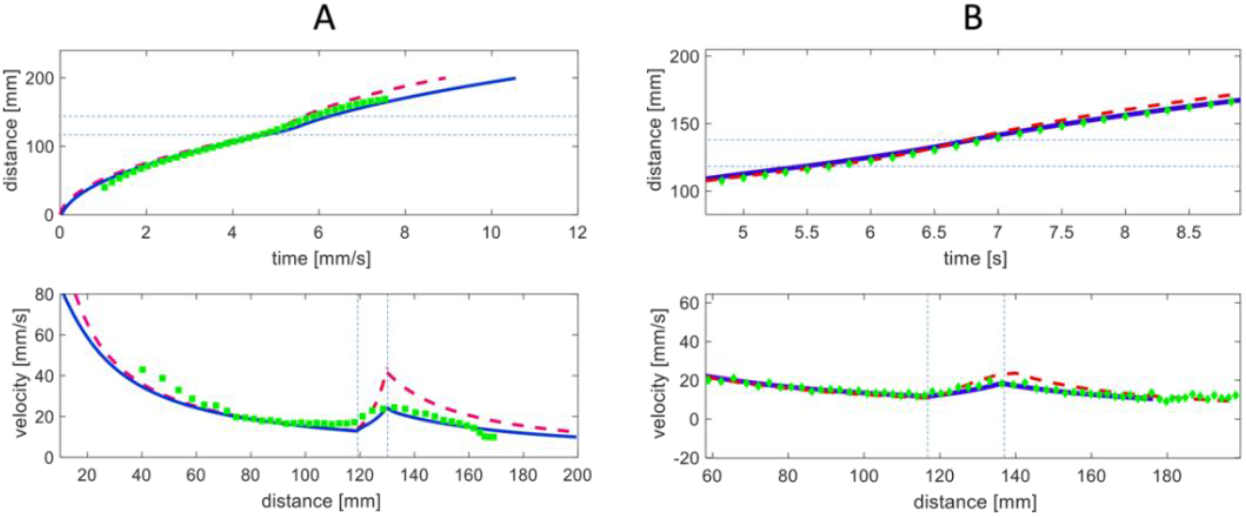
Comparison between experiments and model, of fluid flowing in a converging channel with initial dimensions 2 mm x 2 mm and converging to 1 mm x 2 mm. A: IPA 50% in a device with a 10 mm converging region. B: Pentanol in a device with a 20 mm long convergent region. The horizontal (top) and vertical (bottom) straight dashed lines indicate the boundaries of the convergent region. Top: travel distances vs time; bottom: velocity vs distance. The continuous blue lines correspond to the mBo approach, the green dots to the average of three experiments, and the dashed red lines to the modified LWR approach. Data are reported as one trial (n = 1).

More examples are shown in SI.4, leading to the same conclusions.

## 6. Results for diverging channels

Conversely, the case of divergent channels differs slightly, as previously mentioned in Section 1. In these geometries, no significant inertial effects were observed. This was confirmed through experiments with various geometries and liquids, demonstrating that the inertia-less viscous regime accurately describes the flow dynamics. Figure 11 shows the dynamic of IPA50 flowing in a 20 mm long divergent.

**Figure 11.**
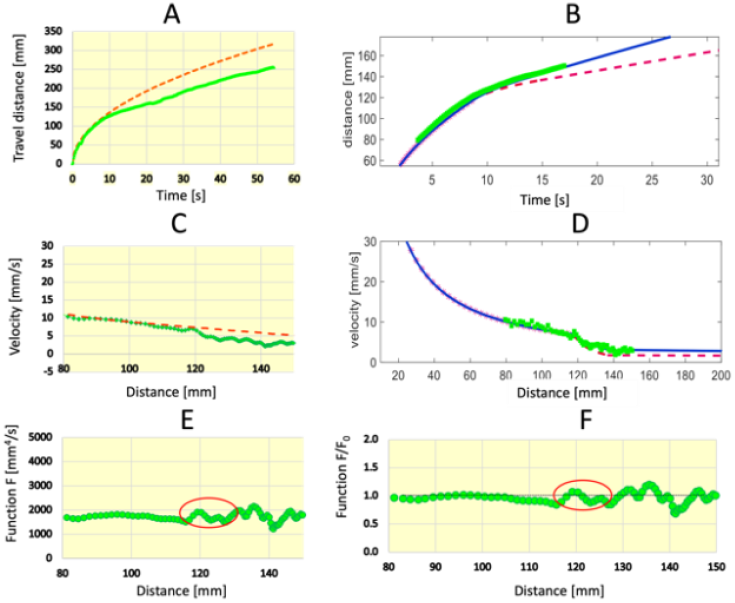
Capillary flow of IPA 50% through a divergent channel (divergent length 20 mm, inlet channel 1 mm x 2 mm, outlet channel 2 mm x 2 mm). A: measured travel distance as a function of time; B: comparison between experiments (green dots), viscous regime (dashed red line), and viscous-inertial model (blue line); C: experimental velocities; D: velocity comparison between experiments (green dots), viscous regime (dashed red line) and viscous-inertial model (blue line); E: the inertia criterion F does not vary substantially in the divergent region; *F*: the non-dimensional function *F/F*_*0*_ is close to unity all along the divergent region. Data is reported as one trial (n =1).

Note that the divergent length must not be too long, nor the divergent angle too steep, as this would lead to the cessation of capillary flow due to a violation of the SCF condition [25].

## 7. Conclusion

This work explores the role of inertial effects in capillary-driven microflows, challenging the widespread assumption that inertia can be ignored in such systems. Traditionally, the LWR law has been applied under the justification that small channel dimensions and high wall friction dominate, overshadowing inertial contributions. However, this study introduces a qualitative criterion, which is backed by a variety of experiments specifically conducted for this research, as well as findings from existing literature. It only involves the monitoring of the advancing fluid meniscus. The results suggest that inertia, although most prominent during the initial stages of flow, can persist under certain conditions, especially in channels with converging geometries. Specifically, in the case of a localized, sudden change in capillary force, the conventional viscous approach remains valid. However, if the capillary force variation extends over a larger region, a viscous-inertial regime emerges. This nuanced understanding could inform the design and analysis of microfluidic systems where inertial effects have been overlooked.

When inertia is significant—when the inertial criterion indicates the presence of a strong inertial effect, as for convergent channels—the use of the Bosanquet equation is required. The theoretical predictions obtained by discretization of the mBo equation were validated through experiments conducted in open rectangular channels made of PMMA. This study shows that the Bosanquet equation of motion provides an accurate model for the dynamics of open capillary flow in rectangular channels. The inertial solution indicates that the liquid velocity increases in a convergent channel, but to a lesser extent than predicted by the non-inertial mLWR solution. On the other hand, no noticeable inertial effect was found in the case of diverging channels.

## Materials and Methods

### Channel design and fabrication

The devices designed for example #1, #2, and the mLWR and Bosanquet analyses features a symmetrical inlet-outlet system about the dynamic region that allows for investigation of both convergent- and divergent-channeled systems. Both the computer-aided design (CAD) and manufacturing (CAM) files were generated using Fusion (Autodesk, San Francisco, California). The channels feature a rounded edge to discourage filament formation. The rounded edge is generated with the use of a round-edged end mill (radius = 0.2413 for the device containing the largest cross-section dimension of 1 mm x 0.5 mm (channel width x channel height) or 0.1905 mm for the device with the largest cross-section dimension of 2 mm x 2 mm) and is not reflected in the CAD file. Devices were milled out of 3.175 mm poly(methyl methacrylate) (PMMA) (#8560K239, McMaster-Carr, Elmhurst, Illinois) using the DATRON neo computer numerical control (CNC) mill (Datron Dynamics, Milford, New Hampshire). The fabrication method for the device in examples #3, #4, and #5 are described in [31], [32], and [34], respectively. Channel dimensions and milling quality were verified using a Keyence wide-area VR-5000 profilometer (Keyence Corporation of America, Itasca, Illinois). Devices were then sonicated in 70% (v/v) aqueous ethanol for 30 min, rinsed in deionized water and left to dry overnight in the fume hood.

### Solvent preparation

For examples #1-2 and the experimental validation using the rectangular open-channel with converging/diverging region, aqueous isopropyl alcohol (IPA) (Fisher Scientific, #A451-4) was used at a mixture of 50% or 20% (v/v). IPA 50% was colored using blue food coloring (McCormick) at a concentration of 12 µL/mL for visualization purposes. Nonanol (Sigma-Aldrich, #131210) and pentanol (Fisher Scientific, #AC16060010) were used for further validation of the model, as liquids with different characteristics. Both nonanol and pentanol were colored with solvent green 3 (Sigma-Aldrich, #211982), solvent yellow 7 (Sigma-Aldrich, #S4016), or oil blue N (Sigma-Aldrich, #391557) at a concentration of 1 mg/mL. The solutions described in examples #3, #4, and #5 are noted in [31], [32], and [34], respectively. For example #5, pentanol with 1 mg/mL of solvent green 3 was used. The aqueous plug was dyed with yellow food coloring (McCormick) for visualization a concentration of 10% (v/v).

### Experimental design

For the experiments using the rectangular open-channel device with the converging/diverging region with the largest channel dimension being 1 mm x 0.5 mm (channel width x channel height), 549 µL of solvent was added to the inlet via a pipet. Once the solvent in the inlet reservoir was depleted, a 64.9 µL refill of solvent was added to the inlet reservoir. For the experiments using the rectangular open-channel device with the converging/diverging region with the largest channel cross-section dimension being 2 mm x 2 mm (channel width x channel height), 2.2 mL of solvent is added into the inlet via a pipet. Once the solvent in the inlet reservoir was depleted, a 300 µL refill of solvent is added to the back of the inlet reservoir. The experimental designs for examples #3 and #4 are noted in [31] and [32], respectively. For example, #5 in Figure 6C to 6E, a 3 µL plug of yellow water was added to the channel approximately 199 mm from the inlet. Afterwards, 950 µL of blue pentanol was added to the reservoir and pipetted slowly to prevent overflow and ensure that the reservoir meniscus remained flat until all of the fluid in the pipet was emptied. Note: images taken in Figure 6A and B were done separately from the results presented in Figure 6C to 6E. Experiments were captured using a Nikon D5300 DSLR high resolution camera at 60 frames per second.

### Data processing

Frames from video recordings were collected at regular intervals (every 10-100 frames, variable for each experiment) using a custom Python code [27]. Screen captures were imported into ImageJ (Fiji, National Institutes of Health, Bethesda, Maryland) and the segmented line tool was used to measure the distance of the liquid front in each frame. Data was transferred to Microsoft Excel where the velocity was extrapolated from the distance vs. time data. MATLAB was also used for data analysis with the mLWR and the Bosanquet models.

## Acknowledgements

This work was funded by NIH grant R35GM128648. The content is solely the responsibility of the authors and does not necessarily represent the official views of the National Institutes of Health. We also acknowledge the Biochemical Diagnostics Foundry for Translational Research supported by the MJ Murdock Trust.

## Data Availability

The data that support the findings of this study are available from the corresponding author upon reasonable request.

