## Supporting Information for "Detection of inertial effects in capillary flows in open and closed channels"

[‡]Co-corresponding authors

*Equal contribution

**Table of Contents**


| **Section** | | **Page** |
|---|---|---|
| SI.1 | The modified Bosanquet approach | S2 |
| SI.2 | The modified Lucas-Washburn-Rideal (mLWR) approach | S5 |
| SI.3 | Engineering drawings of devices | S7 |
| SI.4 | Additional examples of capillary flow in convergent channels | S9 |

#### Section SI.1 – The Bosanquet approach

The open-rectangular channel is a particularly interesting case because these channels are commonly used in microfluidic circuitry. The flow dynamics in such channels have been well-documented when the channel cross-section remains constant. However, the flow dynamics become significantly more complex when the cross-section varies.

In this section, we apply the Bosanquet approach to incorporate the effects of inertia in a linearly converging section, as illustrated in Figure 2.

##### 1. Theoretical approach

It has been shown that the capillary force in uniform monolithic channels can be expressed as

$$F_{cap} = \gamma\, p\, cos\theta^* = \gamma\,\left[\,\left(2h + w_f\right) cos\theta - w_f\right], \tag{SI1.1}$$

where $\gamma$ is the surface tension, *p* the total channel perimeter in a cross section (free and wetted) and $\theta^*$ the generalized Cassie angle. A correction to the contact angle can be done by considering the dynamic contact angle but we do not show it here for simplicity. We just recall that, in the case of open channels, it has been demonstrated that the expression of the dynamic contact angle (DCA), denoted $\theta_d$ , was given by considering the triple contact line friction $\xi$ .

$$cos\theta_d = cos\theta - (\xi/\gamma)V. \tag{SI1.2}$$

Substituting the expression (SI.2) in (SI.1) shows that the capillary force is not constant in a uniform cross-section channel. In the following developments, we use the notation $\theta$ for simplicity instead of $\theta_d$.

In a linearly converging channel with a small geometrical angle $2\alpha$, the capillary force is approximated by

$$F_{cap,conv} = \gamma\,\left[\,2h(\,cos\theta + sin\alpha) - w_f\, cos\alpha\,(1 - cos\theta)\right]. \tag{SI1.3}$$

This expression is derived from Gibbs energy considerations valid for small to moderate velocities. We have extended the use of the formula to this case where there is a small inertial effect. When $\alpha$ = 0, it can be seen that (SI1.3) reduces to (SI1.1).

The wall friction can be determined using the concept of friction length. Consider the friction in the convergent, starting at the abscissa $z_0$. The friction length, denoted as $\bar{\lambda}$, is a purely geometric length that represents the friction within a cross-section, where $\tau = \mu\, V/\bar{\lambda}$. The wall friction is then given by:

$$F_{drag} = \int_{z_0}^{z_f} p\left(\mu\,\frac{V}{\bar{\lambda}}\right) dz = \mu\int_{z_0}^{z_f} p\,\frac{V_f}{\bar{\lambda}}\frac{w_f}{w}\,dz = \mu V_f w_f\int_{z_0}^{z_f}\frac{p}{w\bar{\lambda}}\,dz = \mu V_f w_f G(z_f, z_0). \tag{SI1.4}$$

The integrand $G(z, z_0) = \int_{z_0}^{z_f}\frac{p}{w\bar{\lambda}}\,dz$ is a geometrical value characteristic of the channel. For example, in a rectangular channel of aspect ratio close to 1: $\bar{\lambda} = (w + h)/3(w/h + 2\,h/w)$ and $p = w + h$, hence

$$G(z) = 6\int_{z_0}^{z_f}(1/h + 2h/w^2)dz = 6\left(z_f - z_0\right)/h + 12\, h\int_{z_0}^{z_f} dz/w^2 \ . \quad \text{(SI1.5)}$$

Then, when the flow front is located in the convergent, the Bosanquet equation becomes

$$\frac{d}{dt}\left(w_f\, V_f z_f\right) = \frac{\gamma}{\rho h}\left[\, 2h(\, cos\theta + sin\alpha) - w_f\, cos\alpha\,(1 - cos\theta)\right] - 6\frac{\mu}{\rho h}V_f w_f\int_{z_0}^{z_f}(1/h + 2h/w^2)dz. \quad \text{(SI1.6)}$$

Note that, in the case of a linear convergent, the width $w_f$ is a linear function of the distance, and $V_f = dz_f/dt$. Recalling that the channel width $w$ is linked to the distance $z$ by a linear relation, the last term on the RHS (right hand side) can be integrated and equation (SI1.6) becomes a differential equation in $z_f$:

$$\frac{d}{dt}\left(w_f\,\frac{d}{dt}z_f^2\right) = \frac{2\,\gamma}{\rho h}\left[\, 2h(\, cos\theta + sin\alpha) - w_f\, cos\alpha\,(1 - cos\theta)\right] - 12\frac{\mu}{\rho h}w_f\,\frac{dz_f}{dt}\int_{z_0}^{z_f}(1/h + 2h/w^2)dz. \quad \text{(SI1.7)}$$

Note that the contact angle $\theta$ may depend on the velocity as stated in equation (SI1.2). Under the assumption that the convergent has straight sides, the relation between width $w_f$ and the distance $z_f$ in the convergent region is

$$w_f = w_0 + \frac{(z_f - z_0)}{L_1 - L_0}\,(w_1 - w_0)\,, \quad \text{(SI1.8)}$$

where $L_0$ and $L_1$ are respectively the abscissa of the end of the root channel (before the convergent) and of the end of the convergent (the length of the convergent is then $L_1$- $L_0$). $w_0$ and $w_1$ are respectively the width at the entrance and exit of the convergent. Taking this into account, relation (SI1.8) can be written as

$$w_f = \frac{w_0 L_1 - w_1 L_0}{L_1 - L_0} + z_f\,\frac{(w_1 - w_0)}{L_1 - L_0} = \xi_1 + \xi_2\, z_f. \quad \text{(SI1.9)}$$

Substituting (SI1.9) in (SI1.7) and integrating the last term on the RHS, yields

$$\frac{d}{dt}\left[\left(\xi_1 + \xi_2\, z_f\right)\,\frac{dz_f^2}{dt}\right] = \frac{2\,\gamma}{\rho h}\left[\, 2h(\, cos\theta + sin\alpha) - \left(\xi_1 + \xi_2\, z_f\right) cos\alpha\,(1 - cos\theta)\right]$$
$$-6\frac{\mu}{\rho h}\left(\xi_1 + \xi_2\, z_f\right)\frac{dz_f^2}{dt}\,\left[\frac{1}{h} + \frac{2h}{w_0(\xi_1 + \xi_2\, z_f)}\right]. \quad \text{(SI1.10)}$$

#### 2. Numerical scheme

Let us denote the time step by *Δt*, and we solve equation (SI1.10) iteratively at times {$t_i$, i=1,N} to obtain the travel distances {$z_i$, i=1,N}. The inertial term (noted *I*) can be discretized as

$$I = \frac{d(mV)}{dt} = \frac{\rho h}{\Delta t}\left[(\xi_1 + \xi_2\, z_{i+1})V_{i+1}z_{i+1} - w_i V_i z_i\right], \quad \text{(SI1.11)}$$

where the index *f* has been dropped for simplicity, and $w_i = \xi_1 + \xi_2\, z_i$. On the other hand, the capillary term is

$$F_{cap} = \gamma[2h\ (cos\theta + sin\alpha) - \ w_i cos\alpha\ (1 - cos\theta)]. \qquad \text{(SI1.12)}$$

Finally, the friction term is discretized by

$$F_{drag} = \mu\ w_i V_i \left\{ \frac{p_0\ L_0}{w_0 \bar{\lambda}_0} + 6(z_i - L_0)\ \left\{ \frac{1}{h} + \frac{2h}{w_0\ w_i} \right\} \right\}. \qquad \text{(SI1.13)}$$

Note that the inertial term is discretized for *i+1,* while the values of the two other terms are taken at the time $t_i$. This simplified approach will be proved valid for "fast liquids", such as water, pentanol, IPA50. Combining relations (SI1.11) to (SI1.13) yields

$$\begin{aligned} \frac{\rho h}{\Delta t} [(\xi_1 + \xi_2\ z_{i+1}) V_{i+1} z_{i+1}] = {} & \frac{\rho\ h}{\Delta t} w_i V_i z_i \\ & + \ \gamma[2h\ (cos\theta + sin\alpha) - \ w_i cos\alpha\ (1 - cos\theta)] \\ & - \mu\ w_i V_i \left\{ \frac{p_0\ L_0}{w_0 \bar{\lambda}_0} + 6(z_i - L_0)\ \left\{ \frac{1}{h} + \frac{2h}{w_0\ w_i} \right\} \right\}, \end{aligned} \qquad \text{(SI1.14)}$$

where the unknowns $z_{i+1}$ and $V_{i+1}$ are located at the left hand side. If we use $V_{i+1} = (\ z_{i+1} - \ z_i)/\Delta t$ and regroup the different terms, (SI1.14) becomes

$$\begin{aligned} \frac{\rho\ h}{\Delta t^2} \{ z_{i+1}^3\ \xi_2\ + z_{i+1}^2\ (\xi_1 - \xi_2 z_i) - z_{i+1}\ \xi_1\ z_i \} = {} & \frac{\rho\ h}{\Delta t} w_i V_i z_i \\ & + \ \gamma[2h\ (cos\theta + sin\alpha) - \ w_i cos\alpha\ (1 - cos\theta)] \\ & - \mu\ w_i V_i \left\{ \frac{p_0\ L_0}{w_0 \bar{\lambda}_0} + 6(z_i - L_0)\ \left\{ \frac{1}{h} + \frac{2h}{w_0\ w_i} \right\} \right\}. \end{aligned} \qquad \text{(SI1.15)}$$

Hence, we obtain a polynomial of degree 3 in $z_{i+1}$ .The root can be found using the MATLAB solver and by selecting the "adequate" root (which is the real one, the two others being imaginary roots).

#### Section SI.2 – The modified Lucas-Washburn-Rideal (mLWR) approach

The modified Lucas-Washburn-Rideal (mLWR) model assumes a balance between capillary force (or pressure) and wall friction (or pressure drop due to friction). In the convergent, the balance between the friction force and capillary force is

$$P_{cap} = \frac{p_c\, \gamma cos\, \theta_f^*}{S_c} = \Delta P_f + P_0 = \Delta P_c(z_f) + L_0\, p_0\, \mu \frac{V_0}{\bar{\lambda}_0} \frac{1}{S_0}, \tag{SI2.1}$$

where $\Delta P_f$ is the pressure drop due to friction in the convergent when the tip of the flow is located at the distance $z_f$. Using the relation developed in (SI1.3), the capillary pressure is

$$P_{cap} = \frac{p_f\, \gamma cos\, \theta_f^*}{S_f} = \frac{\gamma\, [\, 2h(\, cos\theta \mp sin\alpha) - w_f\, cos\alpha\, (1 - cos\theta)]}{h\, w_f}, \tag{SI2.2}$$

the "+" sign corresponds to the case of a convergent and the "–" sign to the case of a divergent. The pressure drop can be written

$$\Delta P_c = \int_{z_0}^{z_f} \frac{p}{\bar{\lambda}\, S} \mu\, V\, dz = \int_{z_0}^{z_f} \frac{6\, \left(\frac{w}{h} + \frac{2h}{w}\right)}{w\, h} \mu\, V\, dz, \tag{SI2.3}$$

using $\bar{\lambda} \cong \frac{1}{3} \frac{w+h}{\frac{w}{h} + \frac{2h}{w}}$. Considering the mass conservation equation, the velocity $V$ along the convergent can be expressed in a function of the velocity of the meniscus $V_f$

$$S\, V = S_f V_f, \tag{SI2.4}$$

and (SI2.3) becomes

$$\Delta P_c = \frac{6\mu\, S_f\, V_f}{h^2} \int_{z_0}^{z_f} \frac{\left(\frac{w}{h} + \frac{2h}{w}\right)}{w^2}\, dz. \tag{SI2.5}$$

Let us write $G = \int_{z_0}^{z_f} \frac{\left(\frac{w}{h} + \frac{2h}{w}\right)}{w^2}\, dz$. $G$ is a geometric function of $z_f$ (and $z_0$) and of the convergent/divergent shape that can be written as

$$G(z_f, z_0) = \frac{1}{h} \int_{z_0}^{z_f} \frac{1}{w} dz + \, 2h \int_{z_0}^{z_f} \frac{1}{w^3}\, dz. \tag{SI2.6}$$

In the linear convergent, using the relation developed in (SI1.8) and, after differentiating

$$dw = \frac{(w_1 - w_0)}{L_1 - L_0} dz = -\frac{\delta}{L_c} dz, \tag{SI2.7}$$

where $\delta = w_0 - w_1$ and $L_c = L_1 - L_0$. A change of variables leads to

$$G = \frac{L_c}{\delta} \left\{ \frac{1}{h} \int_{w_f}^{w_0} \frac{1}{w} dw + \, 2h \int_{w_f}^{w_0} \frac{1}{w^3}\, dw \right\}. \tag{SI2.8}$$

Integration of (SI2.8) yields, for a convergent, the function $G^+$

$$G^{+}(w_z) = \frac{L_c}{\delta}\left\{\frac{(\ln w_0 - \ln w_f)}{h} - h\left[\frac{1}{w_0^2} - \frac{1}{w_f^2}\right]\right\}. \quad \text{(SI2.9)}$$

For a divergent where $\delta$ is negative this function becomes $G^{-}$

$$G^{-}(w_{tip}) = -\frac{L_c}{\delta}\left\{\frac{(\ln w_f - \ln w_0)}{h} - h\left[\frac{1}{w_f^2} - \frac{1}{w_0^2}\right]\right\}. \quad \text{(SI2.10)}$$

Using the mass conservation equation and substituting (SI2.9) or (SI2.10), and (SI2.2) in (SI2.1) yields

$$\gamma\,\frac{2h\,(cos\theta \pm sin\alpha) - w_f\,cos\alpha\,(1-cos\theta)}{h\,w_f} = \frac{6\mu\,S_f\,V_f}{h^2}G^{\pm}(w_f) + L_0\,p_0\,\mu\frac{V_f}{\bar{\lambda}_0}\frac{S_f}{S_0^2}. \quad \text{(SI2.11)}$$

Relation (SI2.11) links $V_f$ to $w_f$. Recalling that $V_f = dz_f/dt$, and using relation (SI2.7), we obtain $V_f = -\,(L_c/\delta)\,dw_f/dt$. Substituting in (SI2.11) yields

$$\gamma\,\frac{2h\,(cos\theta \pm sin\alpha) - w_f\,cos\alpha\,(1-cos\theta)}{h\,w_f} = -\mu\,S_f\,\frac{L_c}{\delta}\frac{dw_f}{dt}\left[\frac{6\,G^{\pm}(w_f)}{h^2} + \frac{L_0\,p_0}{\bar{\lambda}_0\,S_0^2}\right]. \quad \text{(SI2.12)}$$

Relation (SI2.12) now links $w_f$ and the time *t*. It can be rewritten as

$$dt = -\frac{h\,w_f}{[2h\,(cos\theta \pm sin\alpha) - w_f\,cos\alpha\,(1-cos\theta)]}\,\frac{\mu}{\gamma}\,S_f\,\frac{L_c}{\delta}dw_f\left[6\frac{G^{\pm}(w_f)}{h^2} + \frac{L_0\,p_0}{\bar{\lambda}_0\,S_0^2}\right], \quad \text{(SI2.13)}$$

which is a relation between the time and the location of the advancing meniscus. Note that $S_f$ is also a function of $w_f$, and (SI2.13) has no closed form solution. But relation (SI2.13) can be numerically solved using the explicit discretization

$$t_{i+1} - t_i = -\frac{h\,w_i}{[2h\,(cos\theta \pm sin\alpha) - w_i\,cos\alpha\,(1-cos\theta)]}\,\frac{\mu}{\gamma}\,S_i\,\frac{L_c}{\delta}(w_{i+1} - w_i)\left[6\frac{G^{\pm}(w_i)}{h^2} + \frac{L_0\,p_0}{\bar{\lambda}_0\,S_0^2}\right], \quad \text{(SI2.14)}$$

where the index *i* corresponds to the preceding (known) time step and *i+1* to the next (unknown) time step. Hence, the numerical approach consists in increasing the travel distance step by step $\{z_i$ , *i=1,n*$\}$, then, using (SI1.8) for determining the width of the channel $\{w_i$ ,*i=1,n*$\}$ and $\{S_i = hw_i\ , i = 1, n\}$ and finally finding the corresponding times $\{t_i$,*i=1,n*$\}$ using (SI2.14).

#### Section SI.3 – Engineering drawings of devices

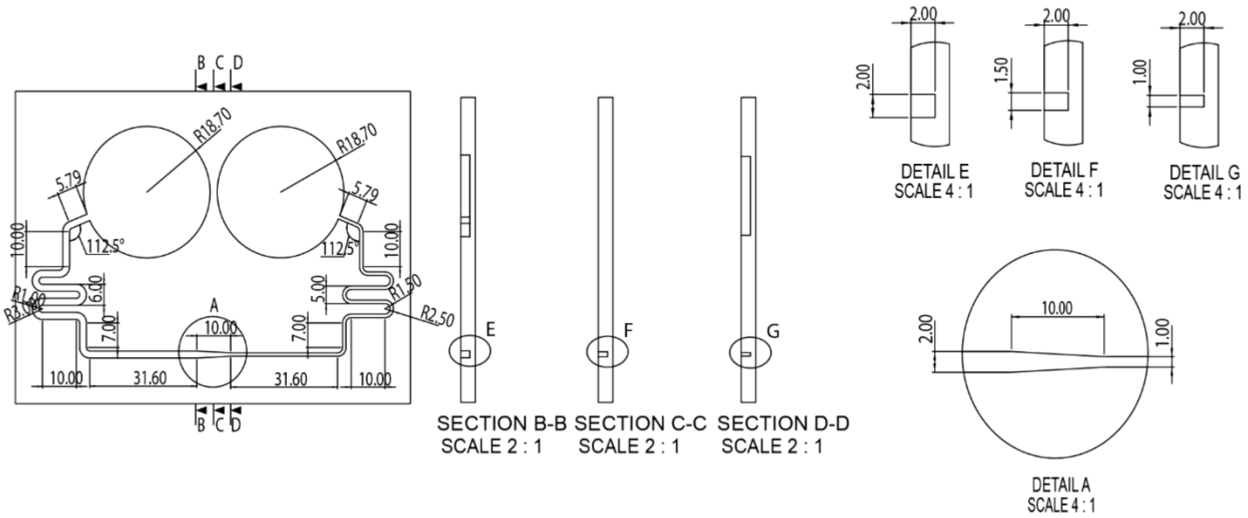


Figure SI.3.1. Engineering drawing of convergent/divergent device with cross-section dimensions of 2 mm x 2 mm reducing to 1 mm x 2 mm over a 10 mm convergent region.

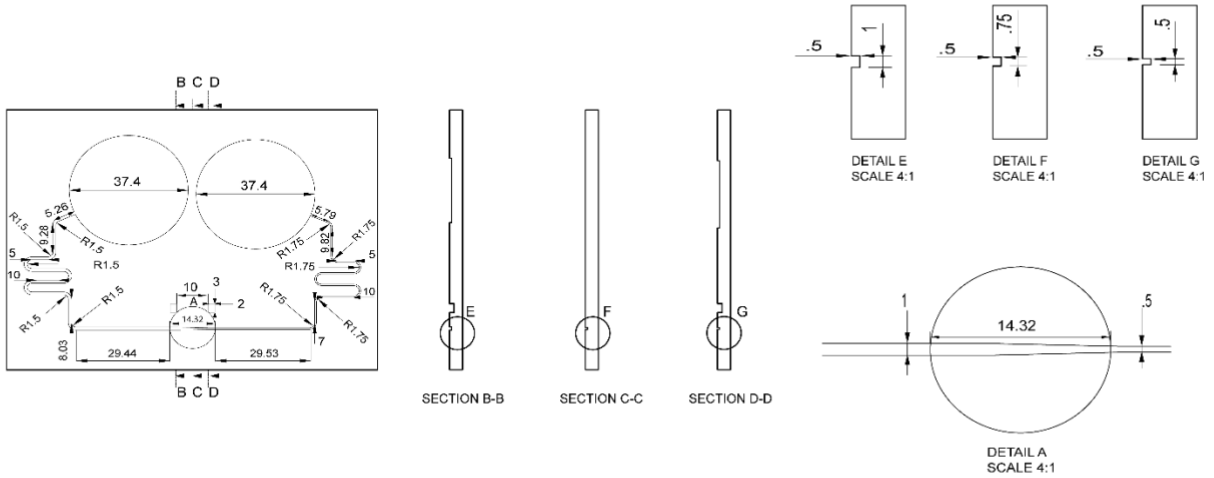


Figure SI.3.2. Engineering drawing of convergent/divergent device with cross-section dimensions of 1 mm x 0.5 mm reducing to 0.5 mm x 0.5 mm over a 10 mm convergent region.

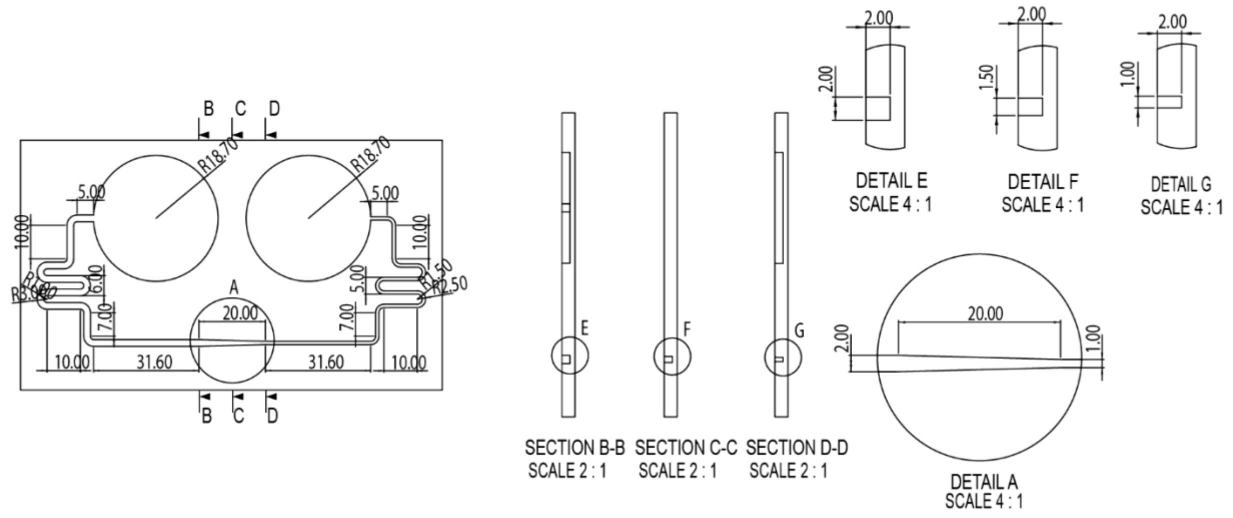


Figure SI.3.3. Engineering drawing of convergent/divergent device with cross-section dimensions of 2 mm x 2 mm reducing to 1 mm x 2 mm over a 20 mm convergent region.

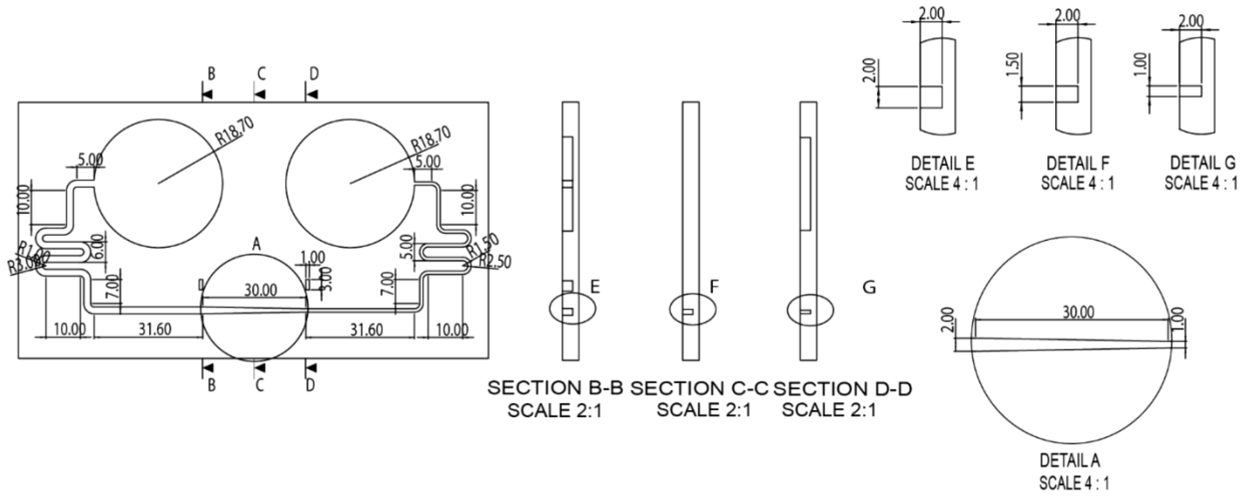


Figure SI.3.4. Engineering drawing of convergent/divergent device with cross-section dimensions of 2 mm x 2 mm reducing to 1 mm x 2 mm over a 30 mm convergent region.

### Section SI.4 – Supplementary examples of capillary flow dynamic in convergent channels

In this SI, we present additional results for the dynamics of a capillary flow  in convergent channels

##### SI4.1. Additional experiments on converging channels

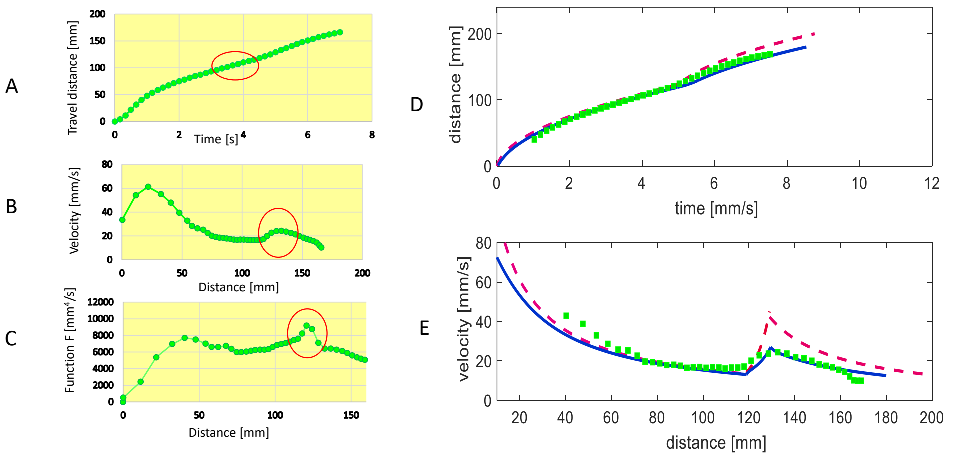


Fig.SI.4.1. Case of IPA 50% v/v flowing in a linear convergent channel of length 10 mm ($w_0$ =2 mm, $h_0$= 2 mm rectangular open channel reducing to $w_1$=1 mm, $h_1$= 2 mm). A: measured travel distances in function of time; B: velocities in function of distances; C: inertial criterion function *F;* D: comparison between experiments (green dots), viscous model (red dashed line) and viscous-inertial model (continuous blue line). The circles in the left pictures indicate the position of the convergent.

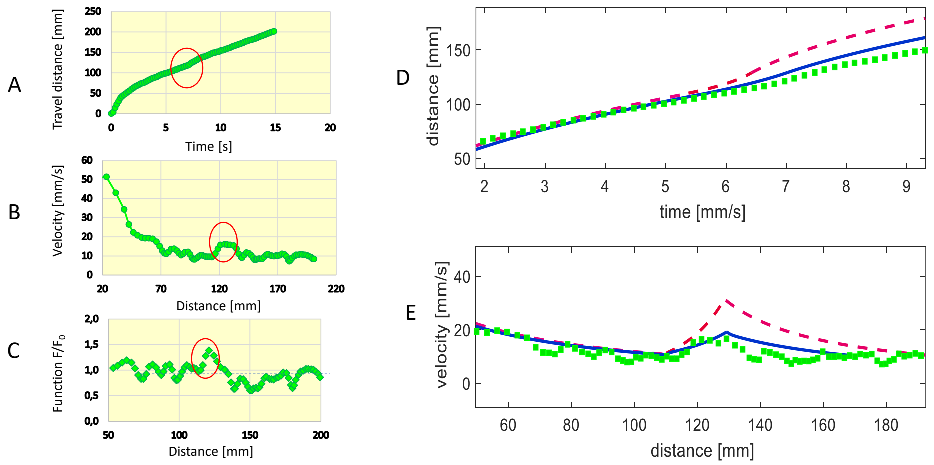


Fig.SI.4.2. Case of IPA 50% v/v flowing in a linear convergent channel of length 20 mm ($w_0$ =2 mm, $h_0$= 2 mm rectangular open channel reducing to $w_1$=1 mm, $h_1$= 2 mm). A: measured travel distances in function of time; B: velocities in function of distances; C: inertial criterion function *F;* D: comparison between experiments (green dots), viscous model (red dashed line) and viscous-inertial model (continuous blue line). The circles in the left pictures indicate the position of the convergent.

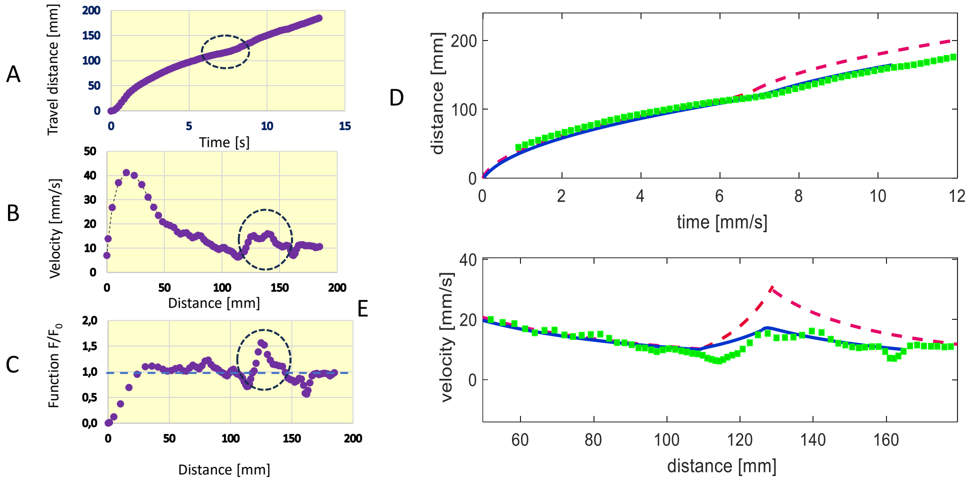


Fig.SI.4.3. Case of IPA 20% v/v flowing in a linear convergent channel of length 20 mm ($w_0$ =2 mm, $h_0$= 2 mm rectangular open channel reducing to $w_1$=1 mm, $h_1$= 2 mm). A: measured travel distances in function of time; B: velocities in function of distances; C: inertial criterion function *F;* D: comparison between experiments (green dots), viscous model (red dashed line) and viscous-inertial model (continuous blue line). The circles in the left pictures indicate the position of the convergent.

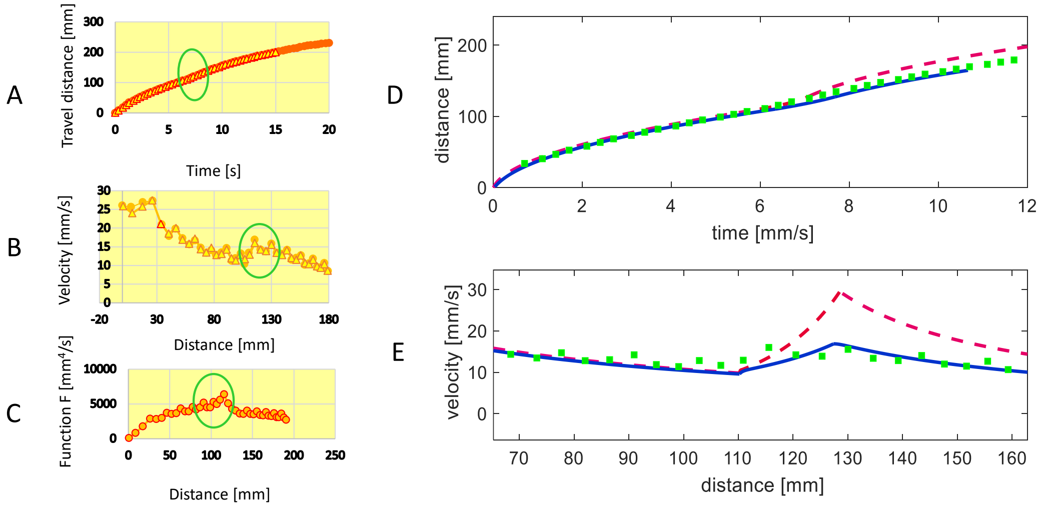


Fig.SI.4.4. Another experiment using IPA 20% v/v flowing in a linear convergent channel of length 20 mm ($w_0$ =2 mm, $h_0$= 2 mm rectangular open channel reducing to $w_1$=1 mm, $h_1$=2 mm). A: measured travel distances in function of time; B: velocities in function of distances; C: inertial criterion function *F;* D: comparison between experiments (green dots), viscous model (red dashed line) and viscous-inertial model (continuous blue line). The circles in the left pictures indicate the position of the convergent.
